# Sequence-based prediction of olfactory receptor responses

**DOI:** 10.1101/664383

**Authors:** Shashank Chepurwar, Abhishek Gupta, Rafi Haddad, Nitin Gupta

## Abstract

Computational prediction of how strongly an olfactory receptor responds to various odors can help in bridging the widening gap between the large number of receptors that have been sequenced and the small number of experiments measuring their responses. Previous efforts in this area have predicted the responses of a receptor to some odors, using the known responses of the same receptor to other odors. Here we present a method to predict the responses of a receptor without any known responses, by using available data about the responses of other conspecific receptors and their sequences. We applied this method to olfactory receptors in insects *Drosophila melanogaster* (both adult and larva) and *Anopheles gambiae*, and to mouse and human olfactory receptors. We found the predictions to be in significant agreement with the experimental measurements. The method also provides clues about the response-determining positions within the receptor sequences.

## Introduction

Odor sensing begins with the binding of odor molecules to olfactory receptors (ORs) expressed on the membranes of olfactory sensory neurons. An organism often expresses a repertoire of tens or hundreds of types of receptors, which combinatorially can detect a very large number of odor stimuli (Malnic et al., 1999). The amino acid sequences of these receptors – and consequently their tuning profiles to odors – vary from species to species, in accordance with their environmental niches.

Identification of the odors that a given OR responds to, also known as deorphanization, is fundamental to understanding olfactory processing. In cases where the sensory neurons expressing a particular OR are known and easy to target with electrodes, the cognate odors can be identified using extracellular recordings from those neurons (Lu et al., 2007; Olsson and Hansson, 2013; Li et al., 2018). An alternative approach is to express the OR in an easy-to-target cell using heterologous expression systems — including cell lines, *Xenopus* oocytes, or the empty neuron system; these have been used in different species including humans (Touhara et al., 1999; Mainland et al., 2015), mice (Oka et al., 2006; Saito et al., 2009), fruit flies (de Bruyne et al., 2001; Hallem and Carlson, 2006; Lin and Potter, 2015), mosquitoes (Carey et al., 2010a; Wang et al., 2010), moths (de Fouchier et al., 2017), and tsetse flies (Chahda et al., 2019). However, such experiments need to be performed separately for each receptor and are time-consuming; moreover, some receptors do not express well in the heterologous systems (Ronderos et al., 2014). High-throughput methods for identifying OR-odor interactions have also been developed recently, but these require elaborate experimental pipelines for each species (Jiang et al., 2015; Hu and Matsunami, 2018; Jones et al., 2019). The high-throughput methods also generate many false positives, so it is recommended that the identified responses be further verified with targeted experiments (Nishizumi and Sakano, 2015).

With lowering costs of sequencing, the sequences of ORs are rapidly becoming available for various organisms, including multiple species of fruit flies (Drosophila 12 Genomes Consortium, 2007; Gardiner et al., 2008; Ramasamy et al., 2016), mosquitoes (Leal et al., 2013; Neafsey and Waterhouse, 2015; Lombardo et al., 2017), bees (Elsik et al., 2015; Karpe et al., 2017) and tsetse flies (Attardo et al., 2019). However experimental data about odor responses are available for relatively few ORs. Computational predictions can potentially bridge this gap. Indeed, computational methods have been developed to predict the response of an OR to new odors based on its known responses to other odors, by analyzing the chemical structure (Schmuker et al., 2007), molecular volume (Saberi and Seyed-Allaei, 2016), or other physicochemical parameters of odor molecules ((Haddad et al., 2008; Boyle et al., 2013; Gabler et al., 2013; Bushdid et al., 2018; Kepchia et al., 2019)). However, these methods can only be used for ORs whose responses to some odors are known.

Here, we present a computational approach for predicting the responses of an OR even when no odor responses are available for that OR; our method instead uses the known responses of other conspecific ORs and the sequence similarities among them. We demonstrate the effectiveness of this approach by comparing our predictions with the experimentally measured responses recordings in insects. Our method also provides insights into the response-determining positions in the receptor sequences.

## Results

### Lack of correlation between OR sequences and responses

As odor responses of an OR depend on its three-dimensional structure, which in turn depends on the amino-acid sequence, we expected that the sequence similarity between a pair of ORs will relate to the similarity in their responses. To test this idea, we used large scale datasets of odor responses of 24 ORs in *D. melanogaster* (Hallem and Carlson, 2006) and 50 ORs in *A. gambiae* (Carey et al., 2010) to sets of 110 odors; these odor sets were partially different between the two species but identical for all ORs within a species. For each pair of ORs within a species, we calculated the distance between their 110-length response vectors (**see Methods**). We quantified amino-acid sequence similarity for each pair of ORs within a species using global sequence alignment. While we expected to see a negative correlation between sequence similarity and response distance, we surprisingly found no correlation at all (**Fig. 1**): the Pearson’s correlation coefficient was 0.0018 (P = 0.97, N = 276 pairs of receptors) in *D. melanogaster*, and 0.026 (P = 0.35, N = 1225 pairs) in *A. gambiae*.

**Fig. 1:**
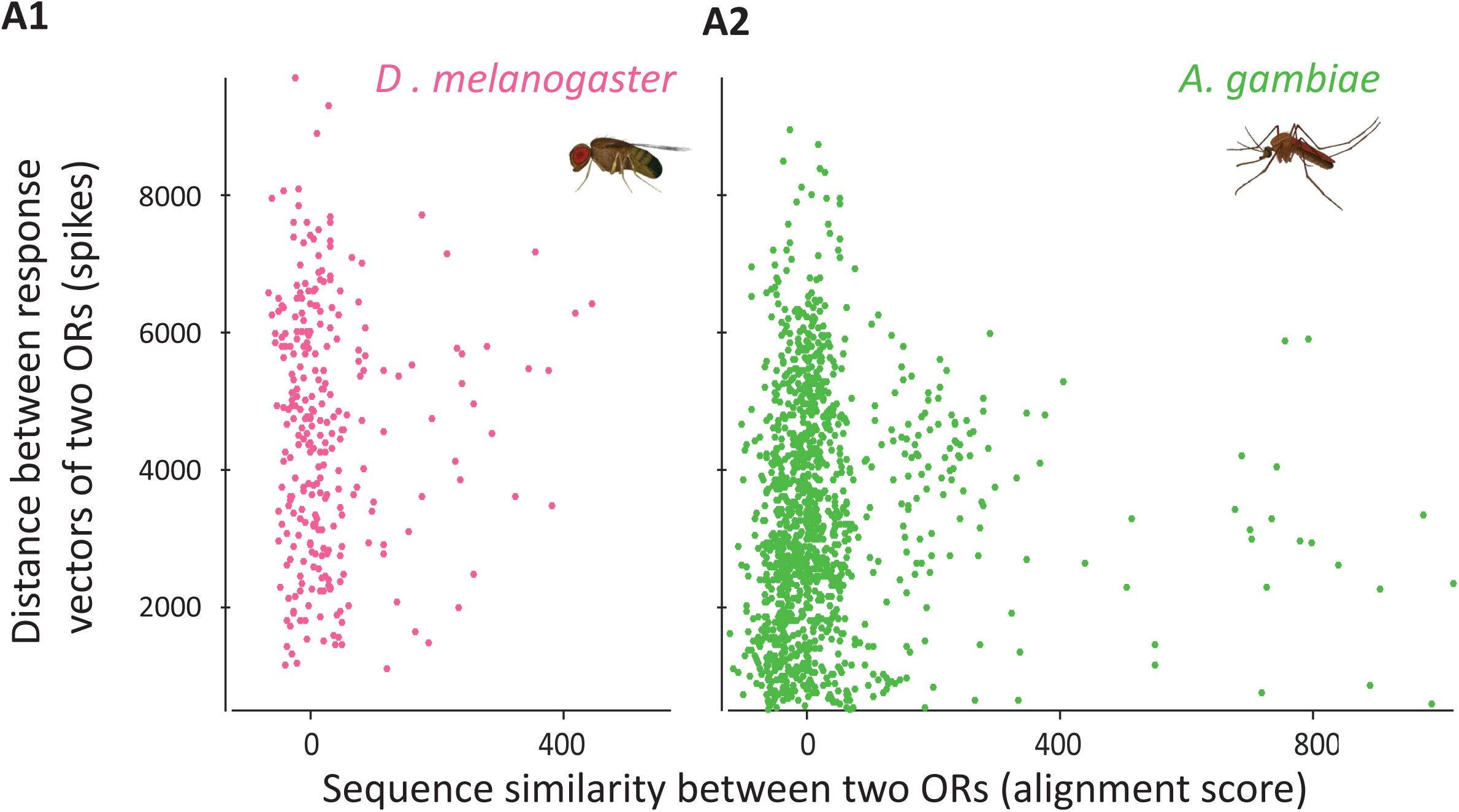
Lack of correlation between response distance and sequence similarity among pairs of ORs. Scatter plots of distances in response vectors versus the amino-acid sequence similarity (a larger value indicates more similarity), among pairs of ORs, reveal the lack of correlation, in both *D. melanogaster* (**A1**) and *A. gambiae* (**A2**). Distance is calculated as the L1-distance between the two 110-length response vectors containing number of spikes for each of the 110 odors. Sequence similarity is measured as the Needleman-Wunsch alignment score. Each point corresponds to a pair of conspecific ORs (n = 276 pairs in *D. melanogaster* and n = 1225 pairs in *A. gambiae*).

### Identification of response-determining positions

The task of sequence-based predictions is made challenging by the very low level of conservation among the ORs: the percentage amino-acid identity between pairs of ORs was only 20.23 ± 4.48 % in *D. melanogaster* and 20.60 ± 8.17 % in *A. gambiae* (mean ± s.d. calculated over all pairs within a species). The lack of correlation between the sequences and the responses is partly because only a small fraction of residues in the sequence are involved in determining odor-specificity, and the similarity of these residues is not represented well by the overall sequence similarity (Man et al., 2007). However, as there is very little structural information available for ORs, the positions of these response-determining residues are not known. We took an empirical approach to rank each position in the multiple sequence alignment of all receptor sequences according to its importance in determining the odor responses (see **Methods**). Within a species, the ranking was done jointly over all ORs to avoid overfitting. Using this ranked list of positions for each species, we selected the top 20 positions that should help in predicting the odor responses of new ORs.

We first confirmed that the subsequences of amino acids formed by these 20 positions in each OR were indeed related to the responses. We found the correlation between the response distance and sequence similarity (calculated using only the subsequences formed by the 20 positions) to be −0.56 (P= 1.7 × 10^−24^, N = 276 pairs; **Fig. 2A1**) in *D. melanogaster*, and −0.39 (P = 1.7 ×10^−45^, N = 1225 pairs; **Fig. 2A2**) in *A. gambiae*. Compared to the result seen with the full sequence (**Fig. 1**), these correlations were significantly negative. Similar negative correlations were observed even if we used only half of the odors for calculating the top 20 positions and the other half for calculating the response similarity (**Supplementary Fig. S1**). If we varied the number of top ranked positions used in the analysis, the correlation coefficients did not change for numbers between 10 and 40 but gradually reduced for larger numbers of positions (**Figs. 2B1,2**), suggesting that lower-ranked residues are not important for the prediction of responses. Subsequently, we used the top 20 positions as the response-determining positions in our analysis (**Figs. 2B1,2** shows that our results would not change if we chose some other number between 50 % and 200 % of this number).

**Fig. 2:**
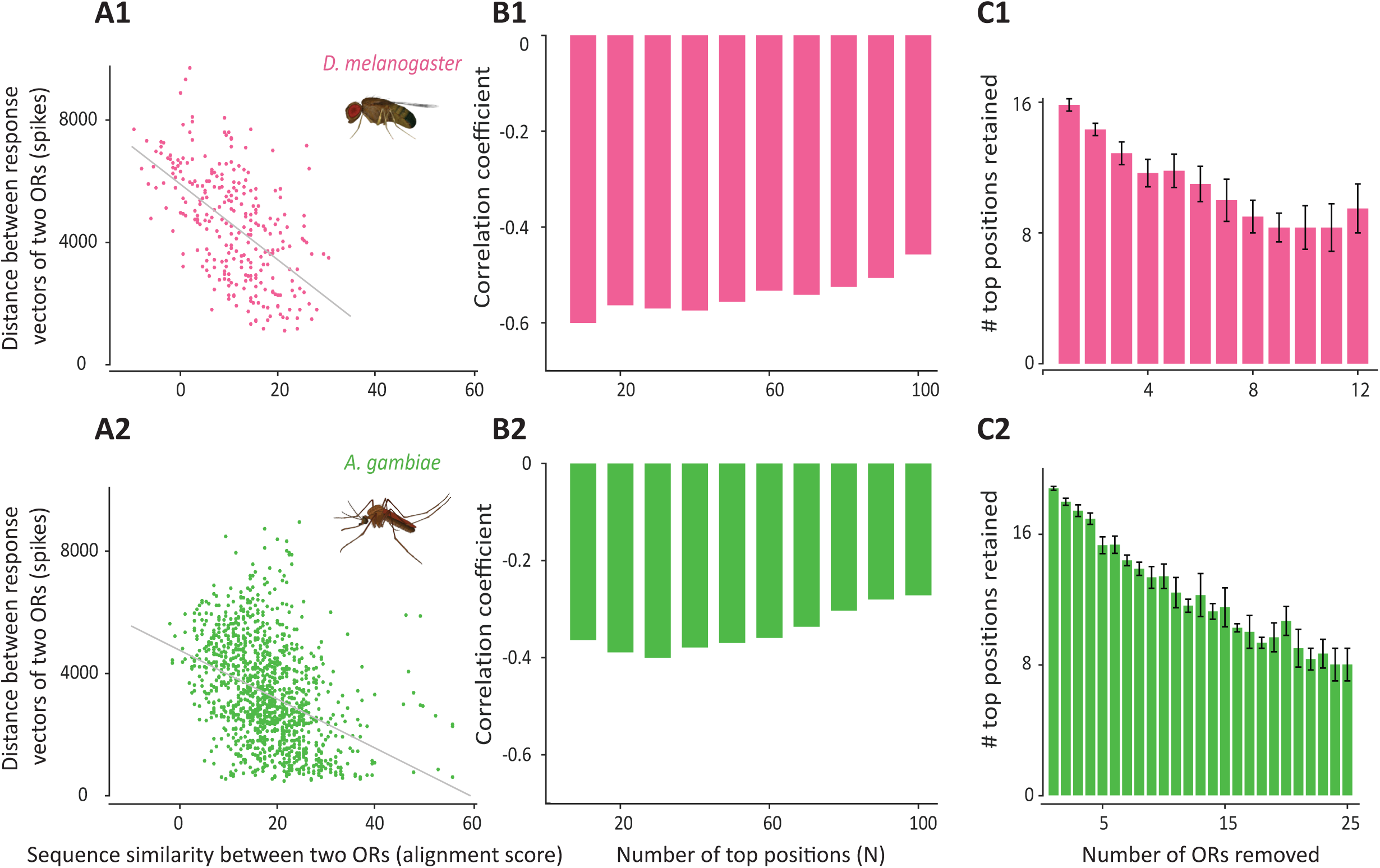
Analysis with response determining positions. **A1, A2**, Scatter plots of distances in 110-odor response vectors versus the sequence similarity measured at the top 20 positions, among pairs of ORs, show significant negative correlation, in both *D. melanogaster* (**A1**) and *A. gambiae* (**A2**). Each point corresponds to a pair of conspecific ORs (n = 276 pairs in *D. melanogaster* and n = 1225 pairs in *A. gambiae*). **B1, B2**, Bars indicate the negative correlation between the distance in response vectors and the sequence similarity measured using the top N positions, plotted as a function of N in the range of 10 to 100. Note that the correlation values are relatively stable in both the species in the range of 10-40 and decrease gradually in magnitude for larger values of N. **C1, C2**, Bars indicate the number of top 20 positions in *D. melanogaster* (**C1**) or *A. gambiae* (**C2**) retained among the top 20 positions when the positions were recalculated after removing different numbers of receptors from the dataset. For each number, all possible combinations of that many receptors were removed and the average number of positions preserved among the top 20 is reported. Error bars represent s.e.m. over all combinations for each number.

We then checked the robustness of the top 20 positions by reducing the number of receptors used for ranking the positions. When we removed one receptor (this was done by removing each receptor, one at a time, to get the averages and error bars), we found that on average ∼16 of the 20 positions in *D. melanogaster* and ∼19 of the 20 positions in *A. gambiae* were retained in the top 20. The number of retained positions dropped gradually as more receptors were left out (again, all possible combinations were tested for each number). But even with half of the receptors removed, at least 8 of the original 20 positions remained in top 20 in both the species (**Figs. 2C1,2**). This analysis indicated that the selected positions were robust to minor perturbations in the datasets.

Although the top 20 alignment positions were chosen independently in *D. melanogaster* and *A. gambiae*, we found that 3 of the top 20 positions were conserved between the species. This number was three times the number of conserved positions expected by chance (20 × 20/369 = 1.08, where 369 is the number of alignment positions that were ranked in both the species), further increasing our confidence in the reliability of the identified positions.

### Predicting responses using the identified positions

Having identified reasonable candidates for response-determining positions, common to all ORs within a species, we asked if it is possible to reliably predict the responses of a query OR to a panel of odors using only the sequence information about the amino acids present at the identified positions. We tested this idea using a simple and intuitive approach, in which the response of a query OR was estimated as the mean of the responses of known ORs, weighted by their similarity to the query OR at the response-determining positions using the BLOSUM62 amino-acid substitution matrix (see **Methods**).

We first applied this approach to predict the responses of the 24 and the 50 ORs in the *D. melanogaster* and *A. gambiae* datasets, respectively. By turn, each OR was treated as a query OR and its response was predicted using the responses of the remaining ORs in the same species (excluding the query OR). We found that the predicted response profiles (i.e., the 110-length vectors of odor responses) of the ORs were correlated with their actual response profiles (**Figs. 3A,B**); the correlations were positive and statistically significant in 21 of the 24 ORs in *D. melanogaster*, and 29 of the 50 ORs in *A. gambiae*. Mean of these correlation values was significantly greater than zero: 0.46 (P = 2.53 × 10^−7^, t-test; P = 1.63 × 10^−5^, sign-rank test; N = 24 ORs) in *D. melanogaster*, and 0.25 (P = 1.18 × 10^−4^, t-test; P = 1.19 × 10^−4^, sign-rank test; N = 50 ORs) in *A. gambiae*. In contrast, control predictions obtained by shuffling the response matrices (see **Methods**) were not correlated with the actual responses (**Fig. 3C**): mean correlation values were 0.0002 (P = 0.95, t-test; N = 24 ORs) in *D. melanogaster*, and 0.0008 (P = 0.65, t-test; N = 50 ORs) in *A. gambiae*.

**Fig. 3:**
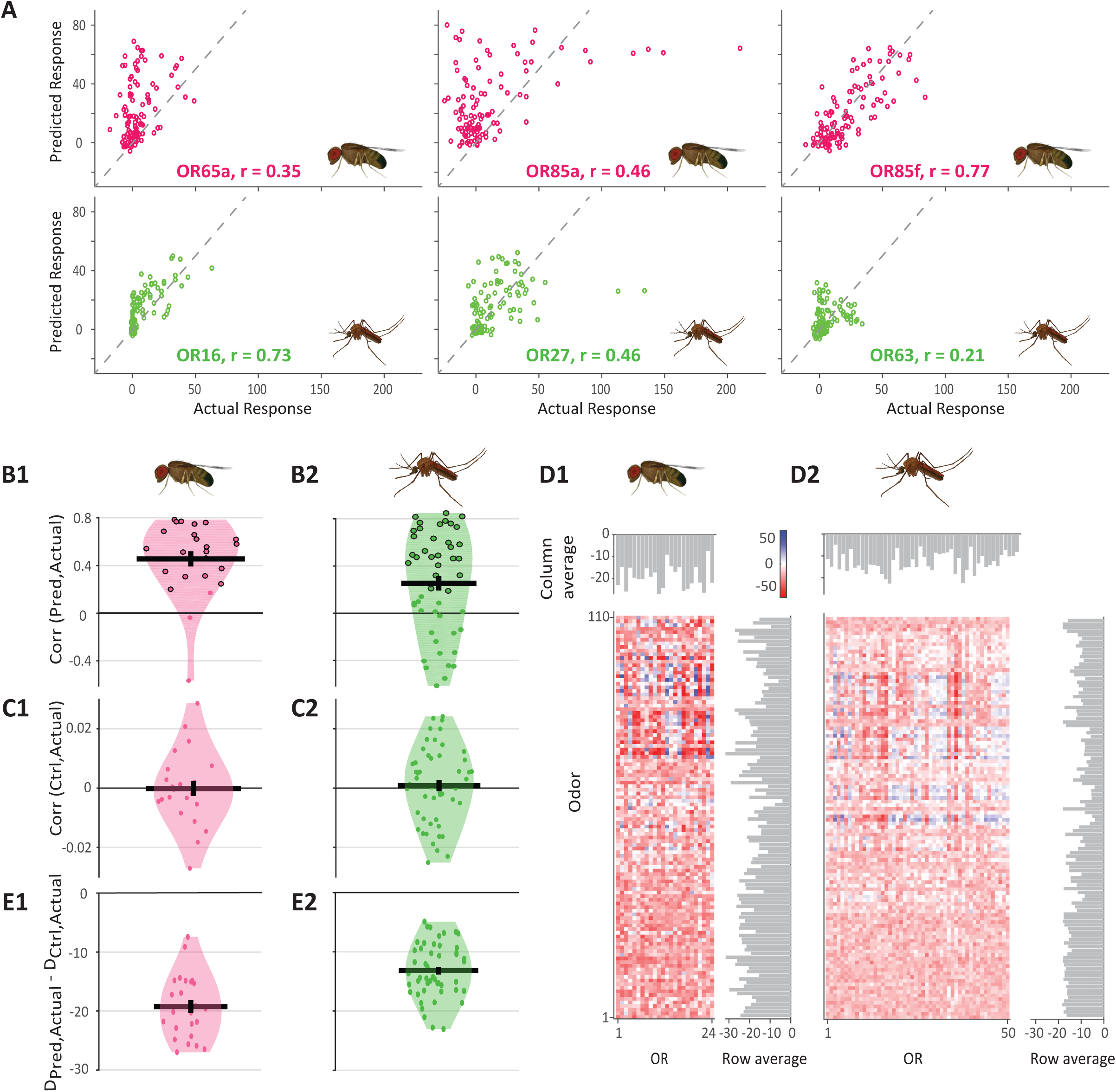
Reliability of responses predicted using the top 20 positions. **A**, Scatter plots of the actual and the predicted odor responses for three representative ORs each from *D. melanogaster* and *A. gambiae*. The correlation coefficients between the two responses are indicated. The responses were spread around the y=x line (dashed line, corresponding to ideal predictions). **B1, B2**, Violin plots showing the correlation coefficient between the predicted and the actual response vectors of ORs in *D. melanogaster* **(B1)** or *A. gambiae* (n = 50) **(B2)**. Each point corresponds to one OR (n = 24 ORs in *D. melanogaster* and n = 50 ORs in *A. gambiae*). Note that the correlations are mostly positive. Statistically significant positive correlations (with P less than 0.05) are shown with black outlines. **C1, C2**, Violin plots showing the correlation coefficient between the control predictions and the actual response vectors of ORs in *D. melanogaster* **(C1)** or *A. gambiae* (n = 50) **(C2)**. Note that the correlations are close to zero in most cases. **D1, D2**, Color maps showing the relative prediction error for each OR-odor pair, for the set of 110 odors, and all 24 ORs in *D. melanogaster* **(D1)** or all 50 ORs in *A. gambiae* **(D2)**. The relative prediction error is measured as *D*_*pred,Actual*_-*D*_*ctrl,Actual*_ where *D*_*pred,Actual*_ is the absolute distance between the predicted and actual response, and *D*_*ctrl,Actual*_ is the absolute distance between the control prediction and the actual response. The abundance of negative values (red shades) indicates that the predictions are usually better than the control. Column averages and row averages of the matrix values are shown on top and right, respectively. **E1, E2**, Violin plots shows the average (over odors) of *D*_*pred,Actual*_ - *D*_*ctrl,Actual*_ for each of the 24 ORs in *D. melanogaster* **(E1)** or 50 ORs in *A. gambiae* **(E2)**. In violin plots, horizontal line and the error bar indicate the mean and the s.e.m., respectively, within a species.

The correlation coefficient between the actual and the predicted odor response vectors tells whether these responses have the same relative magnitude across different odors, but does not tell how close the two responses are in absolute terms (a predicted response that is several times larger than the actual response for every odor will have a perfect correlation). We therefore also used a distance-based metric and checked whether our predictions were closer to the actual responses than the *control* predictions were. We define *D*_*pred,Actual*_ as the absolute difference between the predicted response and the actual response of an OR to an odor, and *D*_*ctrl,Actual*_ as the absolute difference between the control prediction and the actual response. Thus, these two terms indicate the error in the actual and the control predictions. If our predicted response is better (closer to the actual response) than the control prediction, the value of *D*_*pred,Actual*_ − *D*_*ctrl,Actual*_ should be negative. The heat maps in **Fig. 3D** show the comparisons for all the predictions in *D. melanogaster* and *A. gambiae* and reveal a very high abundance of negative values. In *D. melanogaster*, the average *D*_*pred,Actual*_ of an OR (28.60) was smaller than average *D*_*ctrl,Actual*_ (47.84) by 19.24 spikes (P = 7.30 × 10^−15^, paired t-test; N = 24 ORs; **Fig. 3E1**), an improvement of more than 40 %. In *A. gambiae*, the average *D*_*pred,Actual*_ (20.11) was smaller than average *D*_*Ctrl,Actual*_ (33.29) by 13.18 spikes (P = 8.46 × 10^−26^, paired t-test; N = 50 ORs; **Fig. 3E2**), again an improvement of about 40 %. Thus, our predicted responses using the sequence similarity at the response-determining positions were significantly closer to the actual responses than random predictions were. These improvements were substantial considering the absolute values of odor responses in *D. melanogaster* (mean ± s.d. = 32.43 ± 49.29 spikes, N = 24 × 110 OR-odor combinations) and *A. gambiae* (mean ± s.d. = 20.31 ± 37.35 spikes, N = 50 × 110).

### Test on an independent dataset

Next, we collected the responses of 26 other ORs in *D. melanogaster* from 10 different studies (de Bruyne et al., 1999, 2001, 2010; Dobritsa et al., 2003; Goldman et al., 2005; Kreher et al., 2005, 2008; Marshall et al., 2010; Ronderos et al., 2014; Dweck et al., 2015) deposited in the DoOR database (Münch and Galizia, 2016). Each of these ORs had responses available for some of the 110 odors used previously; overall, 68 of the previously used odors were represented in the new dataset. These 26 ORs were entirely different from the 24 ORs used previously in selecting the response-determining positions, and therefore provided an independent test for our approach. Using the set of top 20 response-determining positions as selected earlier from the original *D. melanogaster* dataset, we found that the responses predicted for the 26 novel ORs were significantly better with our approach compared to the control predictions (**Fig. 4A,B**): the average *D*_*pred,Actual*_ of an OR (37.01) was smaller than average *D*_*ctrl,Actual*_ (51.36) by 14.35 spikes (P = 1.13 × 10^−7^, paired t-test; P = 5.96 × 10^−8^, sign-rank test; N = 26 ORs), an improvement of about 28 %. Overall, among the 506 OR-odor responses predicted (non-grey squares in **Fig. 4A**), in 380 (75.1%) cases the predictions were better than the control predictions. These results show that once the response-determining positions have been identified using known responses of some ORs in a species, the responses of other conspecific ORs can be predicted reasonably using only their sequences.

**Fig. 4:**
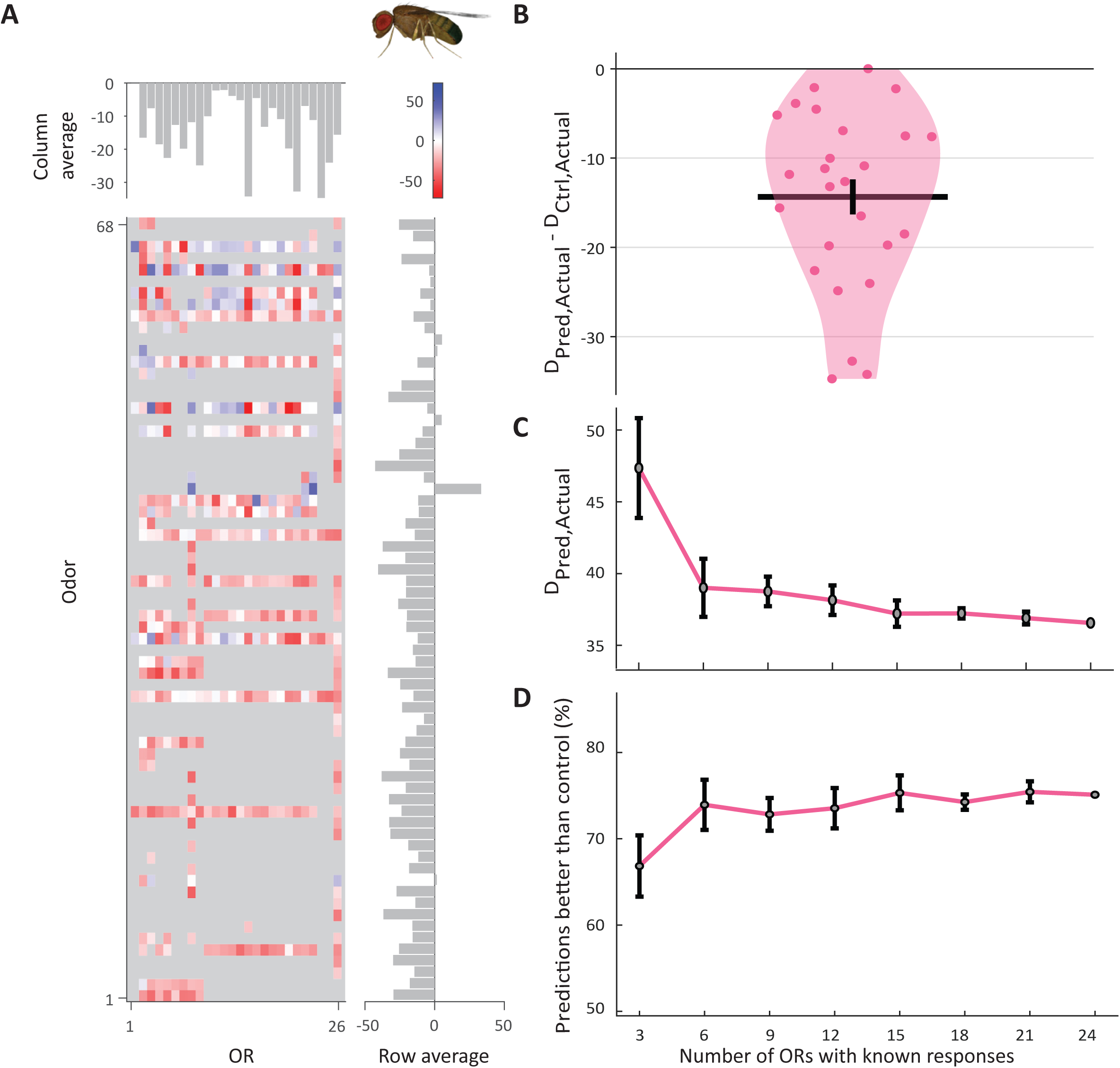
Response prediction on an independent set of ORs. **A**, Color map showing the relative prediction error (measured as *D*_*pred,Actual*_ -_*ctrl,Actual*_) for each OR-odor pair, for the set of 68 odors and 26 novel ORs. Pairs whose values were not available are shown in grey. Note the high frequency of negative values (red) and very low frequency of positive values (blue), indicating that the predictions were closer to the actual value than the control were in most cases. **B**, Violin plots shows the average (over odors) of *D*_*pred,Actual*_-_*ctrl,Actual*_ for each of the 26 ORs. Horizontal line and the error bar indicate the mean and the s.e.m., respectively. **C**, *D*_*pred,Actual*_ is plotted as a function of the number of ORs with known responses used in making the predictions. These different numbers of ORs were sampled from the original set of 24 ORs with known responses. For each number, the sampling was performed 10 times; for each sampling, selection of response-determining positions and prediction of responses was performed. **D**, For each number of sampled ORs, we calculated the fraction of the 506 OR-odor pairs (non-grey values in panel **A**) for which the predicted responses were closer to the actual values than the control predictions were. Both methods of generating control predictions (see **Methods**) were compared. In panels **C** and **D**, errors bars indicate s.e.m. over the 10 samplings for each number from 3 to 21; for 24, all available ORs were used.

While predicting the responses of the new ORs above, we had relied on the known responses of 24 ORs. Can similar predictions be made if responses of fewer ORs are available? To check this, we sampled subsets of ORs from the original dataset of 24 ORs, and used these samples to select response-determining positions and make the response predictions. The sizes of these samples were varied from 3 to 21 in intervals of 3, and for each size the sampling was performed 10 times. We found that the difference between the predicted and the actual response (*D*_*pred,Actual*_) decreased gradually as the number of sampled ORs increased but saturated for 15 or more ORs (**Fig. 4C**). A similar trend was observed when we checked how the number of sampled ORs affected the fraction of predictions that were better than control (**Fig. 4D**). These results suggest that a relatively small set of ORs with known responses may be enough to make reasonable predictions for new ORs.

### Predicting responses in larval *Drosophila*

Although the OR responses in larval *Drosophila melanogaster* differ from the responses in adult *Drosophila* (Kreher et al., 2008), the ORs themselves are the same at the molecular level. Therefore, we checked if the top positions identified in adult *Drosophila* can be used to predict the larval responses. We tested this idea on a dataset of 21 ORs and 26 odors (each available at two different concentrations, 10^−2^ and 10^−4^) from larval *Drosophila* (Kreher et al., 2008). For each concentration, we found that the predictions were significantly better than control predictions (**Supplementary Fig. S2**): at 10^−2^ concentration, the average *D*_*pred,Actual*_ (38.01) was smaller than average *D*_*ctrl,Actual*_ (54.72) by 16.71 spikes (P = 6.32 × 10^−12^, paired t-test; N = 21 ORs); at 10^−4^ concentration, the average *D*_*pred,Actual*_ (12.38) was smaller than average *D*_*ctrl,Actual*_ (16.7) by 4.32 spikes (P = 1.1 × 10^−8^, paired t-test; N = 21 ORs).

### Properties of the response-determining positions

Our results show that the identified subset of sequence positions can allow reliable prediction of the neural responses for new ORs. To check where these top 20 positions are located within the protein structure, we visualized these positions in the secondary structure of two representative ORs in *D. melanogaster* and *A. gambiae* using the Protter tool (Omasits et al., 2014). These residues were not limited to any one region, but were instead found in various parts of the protein (**Fig. 5**). It is important to note that our analysis does not imply a causal or a mechanistic role for each of these residues in interacting with the odor molecules, but only suggests that these residues may be involved in some aspect of determination of odor responses, perhaps indirectly. Notably, the region with the highest density of the top 20 positions was the seventh transmembrane helix, including 4 positions in *D. melanogaster* and 5 in *A. gambiae* (**Fig. 5**; similar numbers were seen in other receptors). Moreover, of the three response-determining positions that were conserved between the two species, two were located in the seventh transmembrane helix. These observations are in agreement with previous studies suggesting that the seventh transmembrane helix plays an important role in determining odor specificity (Hughes et al., 2014; Ray et al., 2014).

**Fig. 5:**
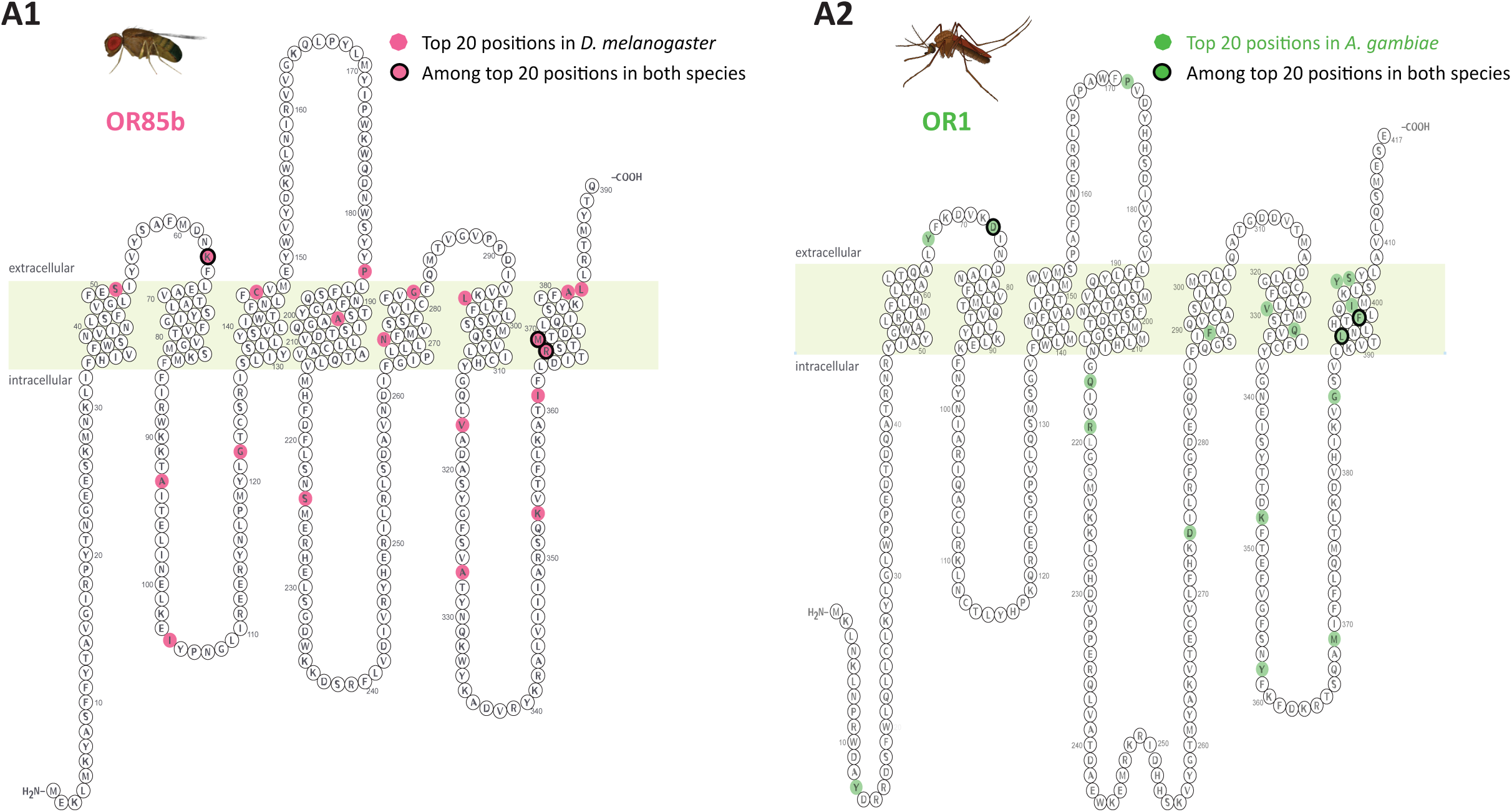
Localization of the top 20 response-determining positions. Protter plots showing the localization of the top 20 response-determining residues in representative ORs: OR85b of *D. melanogaster* **(A1)** and OR1 of *A. gambiae* **(A2)**. Residues in the top 20 positions in the corresponding species are shown with shaded backgrounds, while the residues that are among the top 20 positions in both the species are shown with a black outline. One of the top 20 positions in *A. gambiae* mapped to a gap in OR1 in the multiple sequence alignment, so only 19 positions are shaded.

Next, we collected data from various experimental studies that have mutated individual residues in various ORs and have checked their effect on odor responses. We collected a total of 47 such mutated positions from 5 studies (Nichols and Luetje, 2010; Nakagawa et al., 2012; Xu and Leal, 2013; Hughes et al., 2014; Ray et al., 2014) and compared them with our identified set of response-determining positions. In addition to *D. melanogaster* and *A. gambiae*, these studies also included data from *Bombyx mori* and *Culex quinquefasciatus.* Sequences from these other species were added to our existing multiple sequence alignment (see **Methods**), so that the mutated positions in all these species can be compared with our alignment positions.

Given 20 response-determining positions in *D. melanogaster* and *A. gambiae* each, and with 3 being common between the two species, 37 unique positions in the multiple sequence alignment were marked as the top positions out of a total of 817. Thus, each of the 47 mutated positions in the curated experimental set had a 37/817 probability of matching a top position, and the expected number of matches by chance = 47 × 37/817 = 2.13. So, while only about 2 of the 47 mutated positions were expected to be among the top positions by chance, we found as many as 9 of them to be among the top positions (P = 2.28 × 10^−4^; binomial test), providing experimental support to our response-determining positions. These 9 experimentally studied mutations, which map to the top positions in our alignment, are listed in **Table 1** (further, **Table S1** shows all positions). Among these, residue 146 in OR85b of *D. melanogaster* is one of the residues expected to play a crucial role in the activation of the odorant (Nichols and Luetje, 2010). Residue 109 mutant in OR1 of *B. mori* was found to change the reversal potential and rectification index of the BmOR1-Orco complex (Nakagawa et al., 2012). Point mutation of residues 165 and 194 in BmorOR1 of *B. mori* led to significant reduction in response to bombykol pheromone (Xu and Leal, 2013). Similarly, odor responses in *A. gambiae* were found to change with mutation of residue 369 in OR15 (Hughes et al., 2014), and residues 405 and 406 in OR1 (Ray et al., 2014). Among the remaining 38 out of the 47 mutations in the curated set, 10 and 8 were found to be within a distance of 1 and 2 residues, respectively, from a top column. Thus, in total, 27 of the 47 experimentally confirmed important positions matched or were adjacent to our computationally predicted response-determining positions.

**Table 1:** Positions identified in previous mutagenesis studies that match the top positions identified in our analysis. Of the 47 mutations that we collected from the mutagenesis studies, the 9 that matched the top positions in *D. melanogaster* or *A. gambiae* are listed. The complete set of the 47 mutations is shown in **Table S1**. TM: transmembrane helix; EL: extracellular loop.

| Study | Mutated receptor | Mutated residue | Position in multiple sequence alignment | Rank among top positions | Topological position |
| --- | --- | --- | --- | --- | --- |
| Hughes et al, 2014 | *A. gambiae* OR15 | F369L | 783 | 2 in *D. melanogaster* & 17 in *A. gambiae* | TM 7 |
| Nakagawa et al 2012 | *B. mori* OR1 | Y109A | 234 | 14 in *D. melanogaster* | TM 3 |
| Nichols and Luetje, 2010 | *D. melanogaster* OR85b | C146S | 335 | 20 in *D. melanogaster* | TM 3 |
| Ray et al, 2014 | *A. gambiae* OR1 | Y405H | 791 | 6 in *A. gambiae* | TM 7 |
| Ray et al, 2014 | *A. gambiae* OR1 | S406A | 792 | 20 in *A. gambiae* | TM 7 |
| Xu and Leal, 2013 | *B. mori* OR1 | P165A | 335 | 20 in *D. melanogaster* | TM 3 |
| Xu and Leal, 2013 | *B. mori* OR1 | P194A | 398 | 9 in *A. gambiae* | EL 2 |
| Xu and Leal, 2013 | *A. gambiae* OR10 | P154A | 398 | 9 in *A. gambiae* | EL 2 |
| Xu and Leal, 2013 | *C. quinquefasciatus* OR10 | P157A | 398 | 9 in *A. gambiae* | EL 2 |

### Prediction of vertebrate olfactory responses

As our method does not make any assumption about the composition of the receptor sequences, it should be applicable to any family of chemosensory receptors. We next sought to check its utility in predicting the responses of the vertebrate ORs. Saito et al. (Saito et al., 2009) have provided a dataset of EC50 values of 52 mouse and 10 human ORs that responded to at least one of the 63 odors tested. We extracted a binary response matrix (1 for response, 0 for no response) from this dataset: among the mouse responses (52 × 63 = 3276 values), this matrix included 262 (approximately 8%) 1s and rest were 0s. Using the multiple sequence alignment provided in the same study (Saito et al., 2009) and mouse data, we calculated the top 20 response determining positions. We then calculated the response of each mouse receptor using the weighted average of the responses of the remaining receptors, as done for insect receptors (see **Methods**); for comparison with actual responses, this value was also converted to a binary response using a threshold, set to keep the total fraction of 1s in the predicted responses also at 8%. The predictions thus obtained had a false alarm rate of about 7% and a hit rate of about 21%, resulting in sensitivity index d’ = 0.68 (P < 0.001, estimated by comparing the d’ value with those obtained for 1000 shuffled control predictions). We next asked if human OR responses could also be predicted using the mouse training data. Using the top positions, the responses, and the threshold identified from the mouse data, we predicted the binary responses of the 10 human ORs. Among the 10 × 63 values in the original human dataset, 78 (∼12%) were 1s. Our prediction yielded 581 0’s and 49 1s, with a false alarm rate of about 7% and a hit rate of about 13% (d’ = 0.34, P = 0.047 based on 1000 shuffled predictions). In summary, these results show that our method can also be used to predict vertebrate OR responses with better than chance accuracy.

## Discussion

Sequence-based approaches to response prediction require a reasonably high homology among the receptors. In vertebrates, receptors with less than 40 % sequence identity were found to have little overlap in their responses (Li et al., 2015). The insects ORs presented a particularly difficult test case, as they have only about 20 % homology among conspecific ORs; indeed, we found that there was no correlation between the overall sequence similarity and the response similarity among the ORs. We were able to overcome this difficulty by focusing on subsets of positions that may be important for determining odor responses. Our approach was successful in predicting the responses for not only the set of fly and mosquito receptors whose data was used while developing the method, but also for a completely novel set of receptors whose responses were taken from 10 independent experimental datasets. Further, we found that the predictions require only about 15 ORs with known responses (**Fig. 4**). This computational approach can be particularly useful for receptors expressed in neurons that are difficult to access in electrophysiology experiments, or for receptors that do not express well in heterologous expression systems (Ronderos et al., 2014). Finally, we also showed that our method can also be used for vertebrate receptors.

Our method allows prediction of responses of novel ORs for already studied odors. Previously developed methods allow prediction of responses of already studied ORs to novel odors (Schmuker et al., 2007; Haddad et al., 2008; Boyle et al., 2013; Gabler et al., 2013; Saberi and Seyed-Allaei, 2016; Bushdid et al., 2018; Kepchia et al., 2019). These two approaches are complementary and can be combined in future work to enable the prediction of the responses of novel ORs to novel odors, which would help with large-scale deorphanization. While applying these approaches, care must be taken regarding the concentrations of the odors in the training data, as changes in odor concentration can affect neural responses (Hallem and Carlson, 2006; Olsen et al., 2010) and behavior (Wright et al., 2005; Yarali et al., 2009). Another caveat is that these approaches are currently limited to predicting the response magnitudes, and ignore the temporal patterns of spikes observed in the olfactory receptor neurons (Spors et al., 2006; Raman et al., 2010; Montague et al., 2011; Su et al., 2011; Grillet et al., 2016; Egea-Weiss et al., 2018).

Unlike vertebrate ORs which are G-protein coupled receptors, insect ORs are heteromeric ligand-gated ion channels (Wicher et al., 2008), composed of a specific OR and a conserved co-receptor Orco (Larsson et al., 2004). The structure of any insect receptor was not known until very recently when the structure of Orco from fig wasp was studied using Cryo-EM (Butterwick et al., 2018). The approach of predicting responses using the three-dimensional modeling of ORs, that has seen some success in vertebrates (Bavan et al., 2014), is unlikely to be very productive in insects in the near future given the very low sequence similarity among the ORs and with other proteins, and the paucity of available structures. Therefore sequence-based computational methods will be important for receptors whose structures and responses have not been determined experimentally. The sequence-based predictions are expected to fare better when the responses being predicted are in the same range as the responses in the training dataset, but may not fare as well in predicting uniquely strong ligands for a novel receptor (for the latter goal, three-dimensional modeling may be necessary). Our computational predictions can be used to identify the candidate ligands to be tested experimentally. The method can also be extended to other types of chemosensory receptors, such as the ionotropic receptors and the gustatory receptors.

We have used a simple algorithm in our approach, where we first identify the important positions in the sequences and then use the similarity at those positions to make the predictions. It may be possible to achieve good predictions without explicitly identifying the important positions and instead using a machine learning-based approach that uses all the residues and learns a non-linear mapping between the sequences and the responses. However, we preferred the simpler deterministic approach over the blackbox approaches typically used in machine learning because the deterministic approach provides a transparent rationale for and information about the sequence positions that are used for making the predictions. Expectedly, a majority of the identified positions were located in the extracellular loops or transmembrane helices (particularly the seventh transmembrane, which has been experimentally shown to be important for determining responses), but some were also found in intracellular loops. A previous study has shown that mutation of intracellular residues can affect the electrophysiological responses of neurons (Nakagawa et al., 2012). We emphasize that our approach only looks for correlations between the residues and responses, and it is likely that some of the identified positions, particularly those found in intracellular regions, affect the odor responses indirectly rather than through direct interactions with the odor molecules. By comparing the 47 positions previously found to be important in mutagenesis studies in various organisms (Nichols and Luetje, 2010; Nakagawa et al., 2012; Xu and Leal, 2013; Hughes et al., 2014; Ray et al., 2014), we found that 9 of them (a statistically significant fraction, more than 4 times larger than that expected by chance) mapped to the top positions identified in our analysis (**Table 1**). This shows that our method is using reliable positions for making the predictions. The other top positions that have not been experimentally studied so far are good candidates for future mutagenesis studies, and could help in understanding the mechanisms of odor-OR interactions. As more experimental evidence accumulates, the response predictions can be further improved by giving more weightage to the experimentally verified positions among the top positions.

## Methods

### Response datasets

Responses of 24 *D. melanogaster* ORs were obtained from a previous study (Hallem and Carlson, 2006) and responses of 50 *A. gambiae* ORs were obtained from another study (Carey et al., 2010), for a set of 110 odors. The response values indicate the odor-elicited spiking response of a neuron expressing the receptor, minus the response for the diluent or the spontaneous firing rate. To serve as an independent dataset, responses of 26 other *D. melanogaster* ORs were obtained from 10 studies (de Bruyne et al., 1999, 2001, 2010; Dobritsa et al., 2003; Goldman et al., 2005; Kreher et al., 2005, 2008; Marshall et al., 2010; Ronderos et al., 2014; Dweck et al., 2015) deposited in the Database of Odorant Responses (DoOR) (Münch and Galizia, 2016), for 68 odors that were common with the earlier set of 110 odors. If the response of a particular OR to an odor was reported in multiple studies, we used the average of those values as the actual response.

### Sequence analysis

The olfactory receptor sequences for *D. melanogaster* and *A. gambiae* were obtained from the Database of Olfactory Receptors (Nagarathnam et al., 2014) and the (Hill et al., 2002) study, respectively. The sequence similarity between any two receptor sequences was determined using the Needleman Wunsch Algorithm using the BLOSUM62 scoring matrix (Needleman and Wunsch, 1970; Henikoff and Henikoff, 1992). Multiple sequence alignment was performed for a total of 140 (62 in *D. melanogaster* and 78 in *A. gambiae*) olfactory receptor sequences using MAFFT multiple sequence alignment program, and additional sequences from other species (for validation) were added individually using the ‘-add’ option (Katoh and Frith, 2012).

The OCTOPUS tool (Viklund and Elofsson, 2008) was used to predict the membrane topology of the receptors and the Protter tool was used to visualize the 2-dimensional topology of the receptors (Omasits et al., 2014).

### Ranking the response-determining positions

We identified the response-determining sequence positions for *D. melanogaster* and *A. gambiae* using the following approach, which quantifies at each position how well the differences in amino acids across the receptors correlate with the differences in their responses. These positions corresponded to columns in the multiple sequence alignment, and were same for all the receptors within a species (allowing different positions for each receptor can provide better correlations for individual receptors due to overfitting but will have poor generalizability for other receptors).

From the total of 817 alignment columns in the multiple sequence alignment, we first eliminated the columns which were either fully conserved or had gaps in more than half of the sequences, leaving 378 columns to be ranked in *D. melanogaster* and 392 columns to be ranked in *A. gambiae*, of which 369 were common to both species (see **Table S1**). We denote an element of the alignment by *A*_*r,c*_ where *c* indicates the alignment column and *r* ∈ [1, 24] *or* [1, 50] in *D. melanogaster* and *A. gambiae*, respectively, indicates the row corresponding to a receptor with known responses. The following steps were carried out separately for each species. To calculate the rank for each of the columns, we first calculated a sequence similarity vector (*SS*_*c*_) for each column, each element 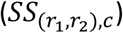 of which contained the pairwise amino acid similarity scores, calculated using BLOSUM62 matrix, between all pairs of receptors (*r*_*1*_,*r*_*2*_)that did not have a gap in the alignment column *c*:

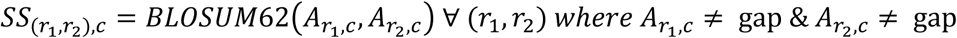

Similarly, we computed a response distance vector 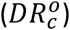 for the same pairs of ORs in the column, each element 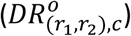 of which contained the absolute difference between the responses of receptors *r*_1_, *and r*_*2*_ to an odor *o*:

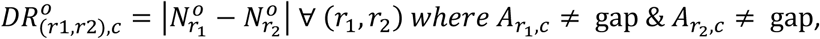

where *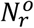* denotes the neural response of receptor *r* to odor *o*. We used the L1-norm instead of the Euclidean distance to avoid giving extra weightage to larger values. Note that *SS*_*c*_ and *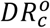* are both vectors of the same length; this length is equal to the number of all possible pairs of receptors that did not have a gap in column *c*. We then calculated the Pearson’s correlation coefficient 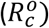 between the sequence similarity (*SS*_*c*_) and response difference 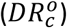 vectors for each column *c* and each odor *o*. The P-value of this correlation, denoted by *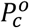*, was also noted. A negative value of *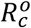* indicates that the receptors that have high sequence similarity at the alignment position *c* also have high response similarity (small distance) for odor *o*. Finally, we calculated the total score of a column as

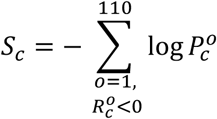

Use of P-values allowed us to give more weightage to highly significant correlations. Columns with high *S*_*c*_ scores were used subsequently as the response-determining positions.

### Response prediction

We first generated the subsequences using the amino acids at the response-determining positions; this was done for the receptors with known responses and also for the receptor whose response is being predicted (*query*). Subsequence similarity (*SSS* _*r,query*_) between the query receptor and each of the known receptors was calculated by taking the average of BLOSUM62 score for all positions in the two subsequences (excluding positions with gaps). The similarity values were then linearly scaled (denoted by 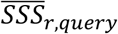) to a range of 0 to 100 to avoid negative weights in the subsequent steps:

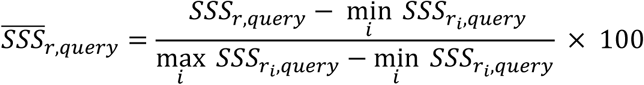

The predicted response 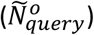 of the query receptor to an odor *o* was calculated as a weighted average of the known responses:

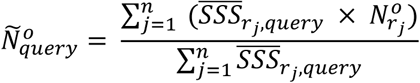

where *n* is the number of receptors with known responses and *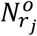* denotes the neural response of receptor *r*_*j*_ to odor *o*.

### Control Predictions

Control predictions were obtained by shuffling the OR-odor response matrix and taking the shuffled response as the control prediction for each OR-odor combination (this operation allowed generation of random predictions while maintaining the overall statistical properties of the response values). The shuffling was not limited to shifting of rows or columns, but included the shuffling of all elements in the matrix. The shuffling was performed 50 times independently, and the results (correlations or *D*_*ctral,Actual*_ values) were averaged over the 50 shufflings.

### Statistical analysis and code availability

All analyses were performed in MATLAB. A modified version of Gramm plotting toolbox (Morel, 2018) was used to draw the plots. We used Pearson correlation to measure the correlations, and t-tests and Wilcoxon signed-rank tests to compare the means. All tests were two-tailed. The number of sample points is shown for each test in the results. The code developed in this study can be accessed from https://github.com/neuralsystems/OR_response_prediction.

## Acknowledgements

We thank members of the N.G. laboratory for helpful comments.

## Author Contributions

N.G. conceived the study. S.C., A.G., R.H., and N.G. developed the methodology; S.C. and A.G. developed the software and performed analysis; N.G. and R.H. secured funding; S.C., A.G., R.H., and N.G. wrote the manuscript.

## Legends

**Supplementary Table S1: Complete multiple sequence alignment of all the ORs.** The table shows the multiple sequence alignment of all the ORs that have been used in this study. Each row corresponds to one OR. The first two rows indicate with colored background the positions that were ranked in the two species; among these the ranks of the top 20 positions are mentioned. The 47 residues that have been identified as important in previous mutagenesis studies are marked with yellow color in the matrix.

**Supplementary Fig. S1:**
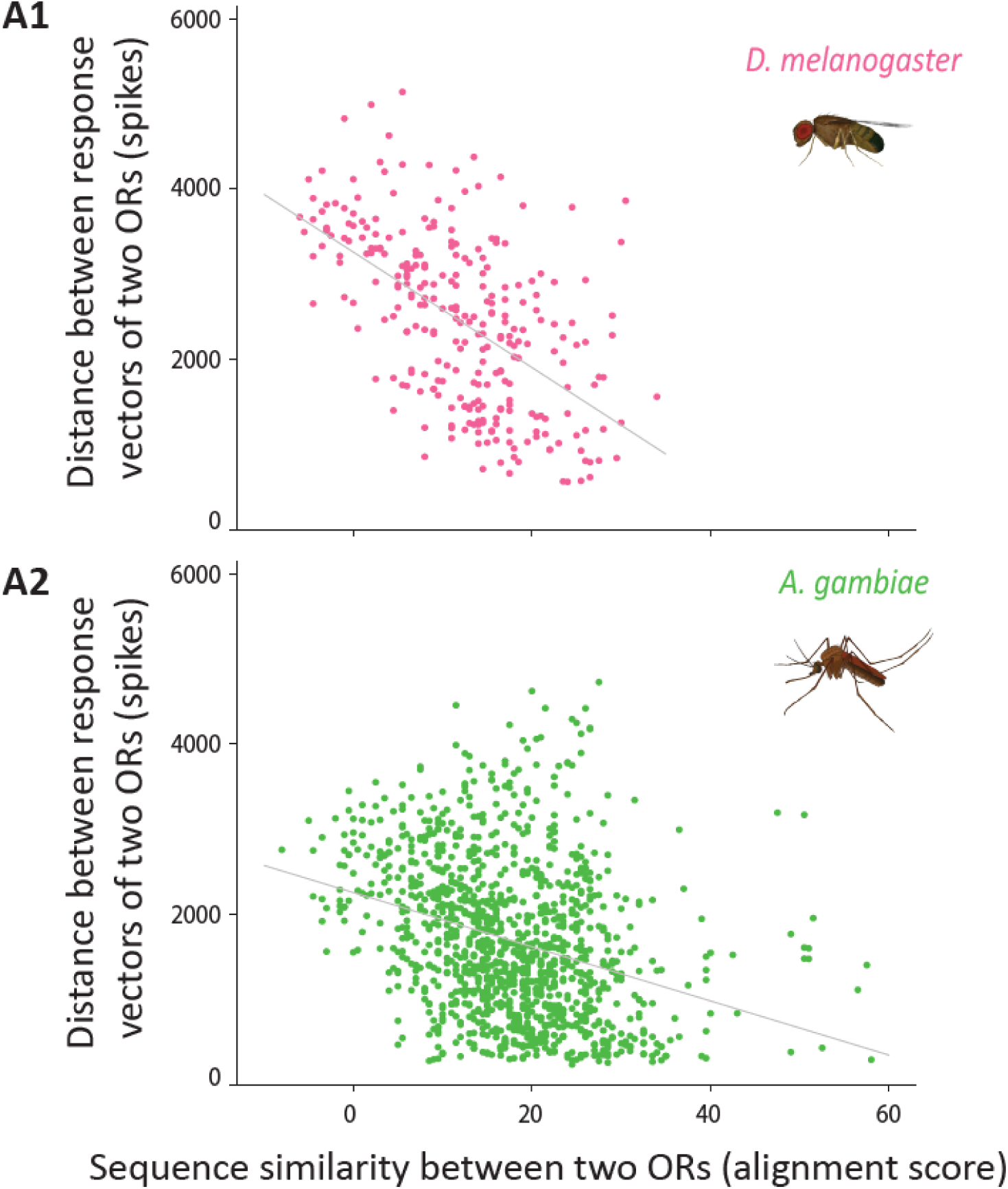
Cross-validation of correlation between response similarity and sequence similarity at the top 20 positions. Scatter plots of distances in response vectors (using half of the odors) versus the sequence similarity measured at the top 20 positions, among pairs of ORs. This figure is similar to Fig. 2A1,2, except that here only half of the odors (in odd-numbered rows of the original dataset) were used to calculate the top 20 positions, and the remaining half were used to calculate the response similarity, thus providing a cross-validation. The observed Pearson correlations are −0.56 in both *D. melanogaster* (**A1**), and −0.31 in *A. gambiae* (**A2**). Each point in the plots corresponds to a pair of conspecific ORs (n = 276 pairs in *D. melanogaster* and n = 1225 pairs in *A. gambiae*).

**Supplementary Fig. S2:**
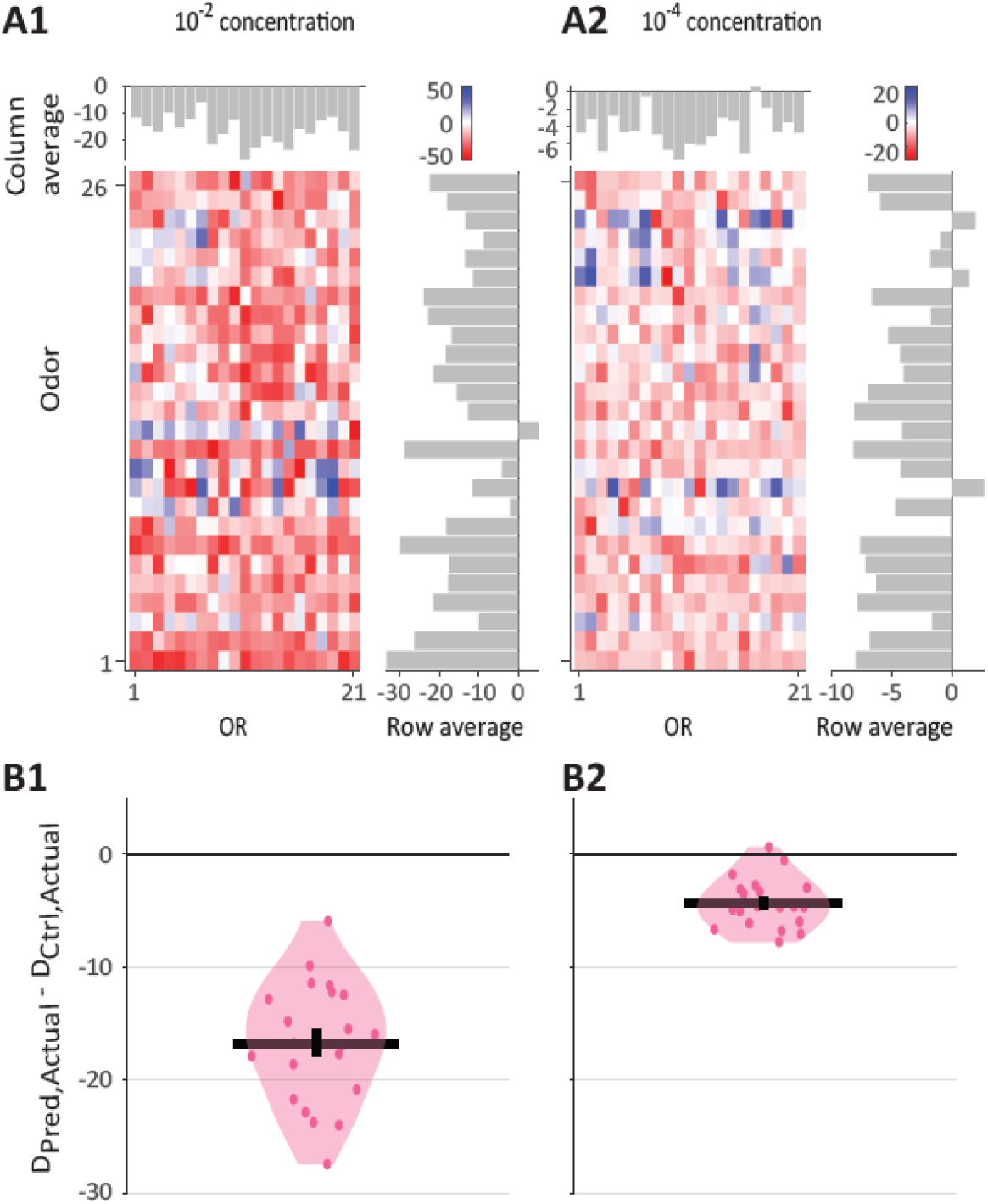
Predictions in larval *Drosophila*. **A1, A2** Color maps showing the relative prediction error for each OR-odor pair, for 21 ORs in *D. melanogaster* larvae for the set of 26 odors, at the dilution of 10^−2^ **(A1)** or 10^−4^ (**A2**). The relative prediction error is measured as *D*_*pred*_,_*Actual*_ - _*ctrl,Actual*_, as described in **Fig. 3.** **B1, B2** Violin plots shows the average (over odors) of *D*_*pred,Actual*_ - _*ctrl,Actual*_ for each of the 21 ORs in panels **A1** and **A2**, respectively. Horizontal line and the error bar indicate the mean and the s.e.m., respectively.

